# A *dwdr45* knock-out *Drosophila* model to decipher the role of autophagy in BPAN

**DOI:** 10.1101/2025.02.06.636873

**Authors:** Marion Celle, Sahra Aniorte, Abdul-Raouf Issa, Marion Falabregue, Haixiu Jin, Irene Sanchez-Mirasierra, Shuzhe Ding, Sandra-Fausia Soukup, Laurent Seugnet, Lujian Liao, Gaetan Lesca, Ludivine Walter, Bertrand Mollereau

**Author notes:** Laboratoire Plasticité du Cerveau, ESPCI-CNRS-UMR8249, INSERM-UA19, Paris, France. INRAE, PEGASE, 35590 Saint-Gilles, France (Marion Falabregue). co-first authors (Marion Celle, Sahra Aniorte).

## Abstract

Beta-propeller protein-associated neurodegeneration (BPAN) is a rare neurological disorder characterized by severe cognitive and motor impairments. BPAN is caused by *de novo* pathogenic variants in the *WDR45* gene on the X chromosome. *WDR45* gene encodes the protein WDR45/WIPI4, a known regulator of autophagy. A defective autophagy has been observed in cellular models of BPAN disease and is associated with neurological dysfunctions in *wdr45* knockout (KO) mice. However, it remains unclear whether the autophagic defect directly contributes to all WDR45 loss-induced phenotypes or whether other WDR45-dependent cellular functions are involved. To investigate this, we generated a CRISPR/Cas9-mediated KO of *CG11975* (*dwdr45* KO), the *Drosophila* homolog of *WDR45*. Our analysis revealed that *dwdr45* KO flies display BPAN-like phenotypes, including impaired locomotor function, seizure-like behavior, autophagy dysregulation and iron dyshomeostasis. Additionally, *dwdr45* KO flies exhibit shortened lifespan compared to control flies. These findings demonstrate that *dwdr45* KO fly is a relevant *in-vivo* model for investigating the key cellular and molecular mechanisms underlying BPAN-associated phenotypes. Here we showed that induction of autophagy in *dwdr45* KO flies improved both the shortened lifespan and the seizure-like behavior, but did not restore locomotor function. This suggests that defective autophagy contributes to some, but not all, aspects of the phenotypes resulting from loss of dWdr45 function.

## Introduction

Beta-propeller protein-associated neurodegeneration (BPAN, MIM 300894) is a rare neurodegenerative disease caused by pathogenic variants in the autophagy-related *WDR45* gene (MIM 300526) (1). BPAN represents one of the most recently identified subtype of neurodegeneration with brain iron accumulation (NBIA), and the increasing number of BPAN diagnoses in recent years positions it as a major subtype, alongside pantothenate kinase-associated neurodegeneration (PKAN). WDR45 is a key regulator of autophagy belonging to the WIPI family (WD repeat domain, phosphoinositide-interacting proteins), which includes WIPI1, WIPI2, WDR45B/WIPI3, and WDR45. These proteins are all involved in autophagy regulation. Most studies have shown that WDR45 primarily regulates bulk autophagy (2). However, findings indicate it also controls selective autophagy, especially ferritinophagy, the process that degrades ferritin to maintain iron homeostasis (3, 4). WDR45 functions at multiple stages of autophagy. It promotes phagophore elongation and controls autophagosome size. Additionally, WDR45 traffics to late endosomes and lysosomes, facilitating autophagosome-lysosome fusion (5, 6). Thus, unlike WIPI1 and WIPI2 which regulate autophagy initiation, WDR45 mainly controls autophagosome maturation and fusion with lysosomes, playing a critical role in advancing the autophagy pathway to completion.

Defective autophagy has been observed in various BPAN cellular and animal models (2, 7–10), although the underlying pathophysiological mechanisms remain poorly understood, and no disease-modifying treatments are currently available for BPAN patients. Thus, there is an urge to develop new experimental models in order to understand the cellular mechanisms underlying BPAN and identify potential cellular targets for therapeutic intervention.

Autophagy is a critical neuroprotective process that is activated in response to cellular stress to preserve neuronal health and lifespan (11, 12). Notably, pharmacological induction of autophagy restored autophagy and alleviated cellular stress in induced pluripotent stem cell (iPSC)-derived dopaminergic neurons from BPAN patients and in *wdr45*-deficient cells, respectively (9, 13). However, it remains unclear whether WDR45 loss-induced autophagy impairment is the main determinant of the BPAN-associated phenotypes.

Iron accumulation in the brain is another hallmark of BPAN and has been extensively studied in cellular models of the disease (14, 15). While iron accumulation has been reported in the brains of aged *wdr45* KO mice, other studies have failed to observe such accumulation (13, 16, 17). Therefore, further investigation using additional animal models of *wdr45* deficiency is needed to study iron homeostasis in BPAN.

*Drosophila melanogaster* is a powerful genetic model organism for studying neurological diseases, including Parkinson’s, Huntington’s, and Alzheimer’s diseases (18). It is particularly well-suited for genetic studies due to its low genetic redundancy and the presence of functional orthologs/homologs for 60% to 70% of genes implicated in human diseases (19). Moreover, autophagy and iron homeostasis mechanisms are largely conserved between flies and mammals, with key components such as ferritin proteins playing central roles in iron storage and detoxification (20, 21).

In this study, we generated and characterized a novel *dwdr45* KO in *Drosophila* to investigate the cellular and organismal phenotypes associated with dWdr45 loss of function. Our findings reveal multiple phenotypic manifestations reminiscent of BPAN, including impaired autophagy and iron dyshomeostasis at the cellular level, as well as motor deficits, seizure-like behavior and reduced lifespan at the organismal level. Furthermore, we demonstrate that genetic neuronal induction of autophagy in *dwdr45* KO flies rescues the shortened lifespan and seizure-like behavior, but does not restore locomotor dysfunction, suggesting that dWdr45-mediated autophagy contributes to some, but not all, of the phenotypes associated with dWdr45 loss of function. This newly established BPAN fly model thus provides a valuable *in vivo* model to dissect the molecular mechanisms underlying BPAN and to evaluate potential therapeutic strategies.

## Results

### Identification of putative WDR45 homologue in *Drosophila*

We first conducted a phylogenetic analysis comparing *Homo sapiens* WDR45/WIPI4 protein sequence with its *Homo sapiens* paralog WDR45B/WIPI3 and with its orthologs from *Mus musculus, Drosophila melanogaster, Caenorhabditis elegans, and Saccharomyces cerevisiae* (**Figure 1A**). Sequence alignment of WDR45 revealed *CG11975* as the putative *Drosophila* homolog of *WDR45*, hereafter referred to as *dwdr45*. Both the fly and human WDR45 share a conserved seven-blade beta-propeller, structure typical of WIPI proteins, along with conserved ATG2 interaction sites and a phosphoinositide-binding domain, known to interact with autophagy regulators (**Figure 1B, C**).

**Figure 1.**
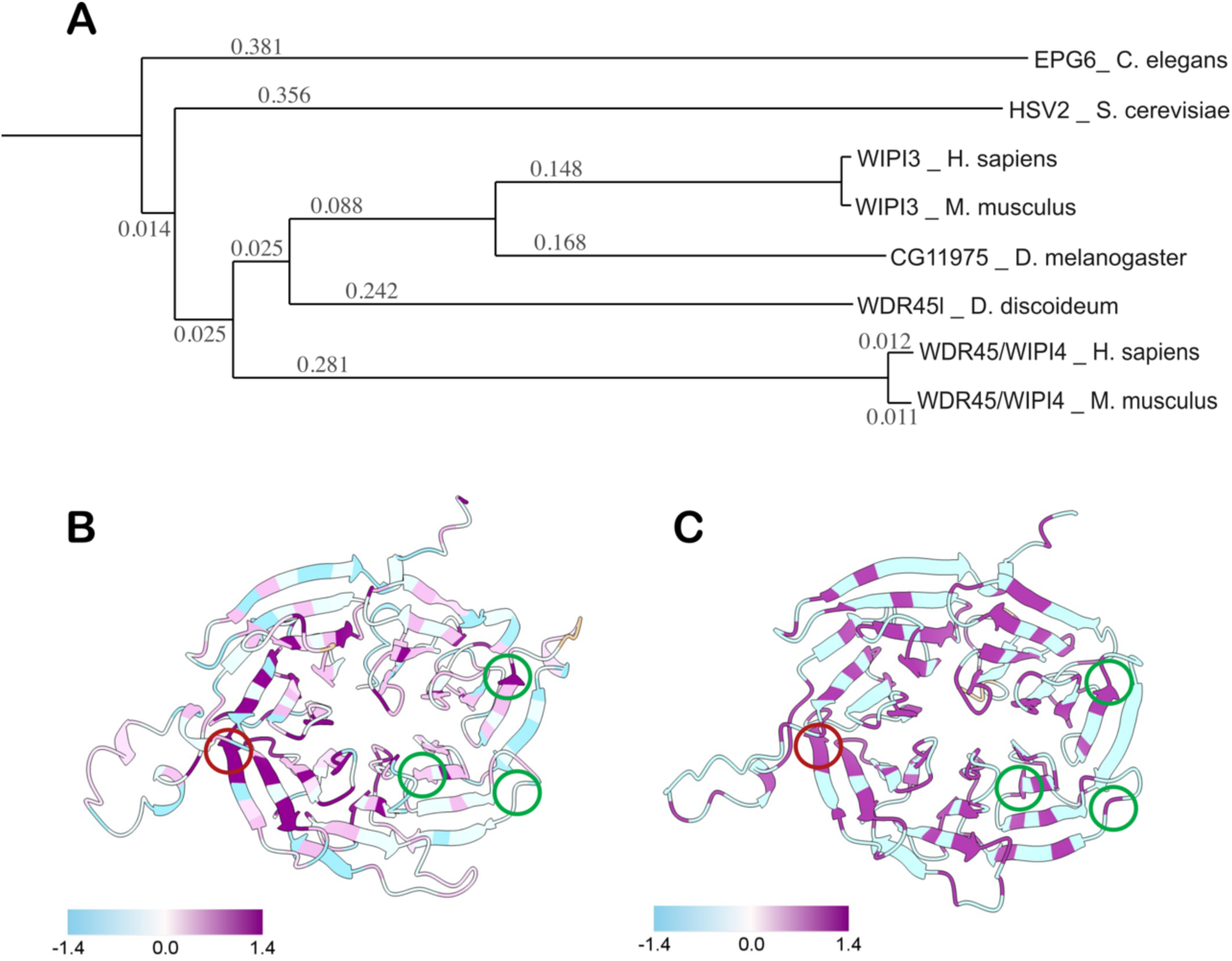
CG11975/dWdr45 is the *Drosophila* homolog of WDR45. (**A**) Phylogenetic tree based on amino acids analysis of hWDR45, its paralog in *Homo sapiens*, and its orthologs *Mus musculus*, *Drosophila melanogaster*, *Dictyostelium discoideum* and *Saccharomyces cerevisiae*. Branch’s length represents evolutive distance. (**B**) Chimeric representation of hWDR45 protein colorizing on conservation index with its orthologs. Phosphoinositide binding domain and ATG2 binding sites are highlighted with red and green circles, respectively. (**C**) Chimeric representation of dWdr45 colorizing on conservation index with hWDR45. Phosphoinositide binding domain and ATG2 binding sites are highlighted with red and green circles, respectively.

Given that *wdr45* pathogenic variants are associated with severe neurological impairments in BPAN (2), we sought to examine the neuronal expression of *dWdr45* in the adult *Drosophila* brain. Using a knock-in GAL4 line, Kozak-GAL4 CG11975^KO-kG4^ promoter trap line combined with UAS-GFP, we tracked *dWdr45* cellular expression through the GFP labelling, and observed a widespread expression of *dWdr45* throughout the adult fly brain, with particularly strong expression in neurons marked by the anti-Elav antibody (**Figure S1**). The neuronal expression of *dWdr45* is also supported by the SCOPE single-cell sequencing data set (22, 23).

### Neuronal, locomotor and seizure-like phenotypes in *dwdr45* KO *Drosophila*

To investigate the cellular consequences of *dwdr45* deficiency, we generated a *dwdr45* CRISPR/Cas9-mediated knockout (KO) predicted to produce a peptide of 38 amino acids in length, in contrast to the 340 amino acids of the full-length wild-type protein. This truncated peptide lacks the characteristic seven-bladed beta-propeller and the autophagy regulator interaction domains (**Figure S2**). Therefore, the *dwdr45* KO flies are likely to carry null allele. BPAN is accompanied by locomotor dysfunction and epilepsy in Humans. To determine whether *dwdr45* KO flies recapitulate these key features of BPAN, we first assessed their locomotor abilities using the startle-induced negative geotaxis (SING) assay. This assay is a well-established tool to measure motor skills and neuronal integrity in *Drosophila* disease models, including Parkinson’s disease ((24, 25) and **Figure 2A)**. Both homozygous *dwdr45* KO flies or transheterozygous between *dwdr45* KO and the GAL4 Kozak insertion in *dwdr45* (*dwdr45^KO^/CG11975^KO-kG4^*), exhibited a reduced performing index compared to control flies, with a further decline as the flies aged, indicating progressive locomotor dysfunction, a well characterized feature of BPAN (**Figure 2B, S3A**). Importantly, neuronal and endogenous re-expression of *dWdr45* rescued the locomotor defects in *dwdr45* KO homozygous and in transheterozygous *dwdr45 ^KO^/CG11975^KO-kG4^* flies (respectively **Figure 2C, S3A**). It indicates that the locomotor deficit is caused by *dWdr45* loss and that *dwdr45* expression in neurons is necessary for proper locomotor function in flies. In addition, we observed similar rescue effect of the locomotion phenotype with an expression of human *WDR45* (*hWDR45*) in the neurons of *dwdr45* KO flies (**Figure 2C**), supporting the phylogenetic analysis (**Figure 1**) and the assumption that dWdr45 is the *Drosophila* functional homolog of hWDR45. Furthermore, knockdown of *dwdr45* using RNA interference (RNAi) under the *Elav* driver also resulted in progressive locomotor deficits similar to those provoked by alpha-Synuclein overexpression, a known motor deficit inducer (**Figure S3B**). Collectively, these results establish that decreased or loss of dWdr45 function cause progressive locomotor dysfunction in flies.

**Figure 2.**
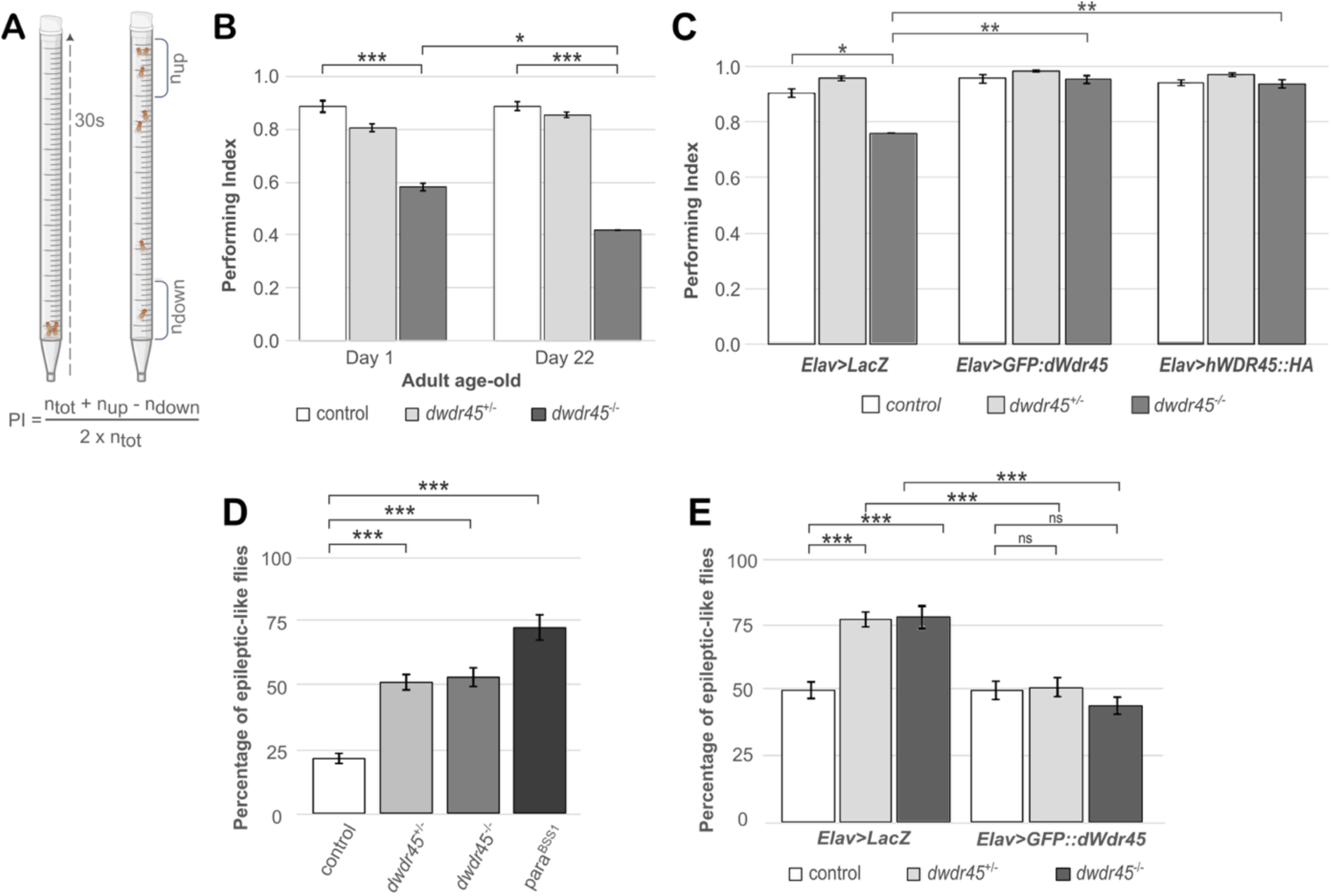
Neuronal overexpression of *dWdr45* rescued locomotor defects and seizure-like phenotypes in dwdr45 KO flies. **(A)** Schematic representation of the SING assay and formula used to calculate the performing index (see Materials and Methods section). **(B)** Representative experiment of three SING assays (n=3) realized in 1- or 22-day-old adult control, heterozygous d*wdr45* KO or homozygous *dwdr45* KO flies. Statistical analyses were carried out with ANOVA2/post-hoc test: *p- value<0.05, **p-value<0.01 and ***p-value<0.001. **(C)** SING assays were realized in 8-day-old adult control (*Elav-Gal4;;*), heterozygous *dwdr45* KO or homozygous *dwdr45* KO flies (*Elav-Gal4;;dwdr45^KO^*) expressing *LacZ*, *GFP::dWdr45* or human *hWDR45::HA* with the pan-neuronal *Elav* driver. Statistical analyses were carried out in three independent experiments (n=3) with ANOVA2/post-hoc test: *p-value<0.05 and **p-value<0.01. **(D)** Percentage of flies exhibiting seizure-like phenotype, described as loss of posture, leg shaking, random wing flapping, spinning and uncontrolled flight, was measured in 4-day-old control, heterozygous *dwdr45* KO or homozygous *dwdr45* KO flies or *para^BSS1^* epileptic fly mutants. Statistical analyses were carried out in three independent experiments (n=3) with ANOVA1/post-hoc test: ***p-value<0.001. **(E)** Percentage of flies exhibiting seizure-like phenotype in control (*Elav-Gal4;;*), heterozygous *dwdr45* KO or homozygous *dwdr45* KO flies (*Elav-Gal4;;dwdr45^KO^/TM6B,Tb*) either expressing *LacZ* or overexpressing *GFP::dWdr45* under the pan-neuronal *Elav* driver. Statistical analyses were carried out on three independent experiments (n=3) with ANOVA1/post-hoc test: ***p-value<0.001 and ns no significant differences.

We next examine whether *dwdr45* KO flies display a seizure-like phenotype. To induce seizures, we applied mechanical shaking, a method classically used to provoke seizure activity in well-characterized *Drosophila* epilepsy models, such as the *para^Bss^* mutant (*para^Bss1^*), which is highly sensitive to mechanical stimulation and exhibits robust seizure-like behavior (26, 27). The proportion of flies displaying abnormal postures such as being on their backs, wing buzzing, and leg shaking was monitored as an indicator of seizure activity. We found that both heterozygous and homozygous *dwdr45* KO flies exhibited seizure-like behaviour comparable to that observed in the *para^Bss1^* mutant in the assay (**Figure 2D**). Moreover, pan-neuronal expression of *dwdr45* under the Elav driver rescued the seizure-like phenotype in *dwdr45 KO*, suggesting neuronal origin (**Figure 2E**).

In BPAN mouse models, neurodegeneration phenotypes range from neuronal loss in specific areas to more subtle neuronal structural defects, depending of the mutant and genetic background considered (13, 16, 17, 28). Given that a locomotor decline is often associated with a loss of dopaminergic neurons across different species including flies (29), we first visualized various clusters of these neurons (PAL, PPL1, PPL2, PPM1/2, and PPM3) using TH staining and we did not observe TH staining changes in the adult brain of *dwdr45* KO flies when compared to control flies (**Figures 3A**), suggesting that *dwdr45* KO flies present no neuronal loss. Although, we cannot exclude the presence of structural neuronal phenotypes in specific neuronal subpopulations of adult or aged brains, or during nervous system development. To test this latest hypothesis, we took advantage of the embryonic nervous system and larval neuromuscular junction (NMJ) in *Drosophila* that are well-defined, accessible, and amenable to imaging techniques (30). These analyses facilitate detailed visualization of individual neurons, axons, dendrites, and synaptic structures, allowing detection of subtle morphological changes that may not be apparent in the adult brain. We first found that the *dwdr45* KO embryo exhibited increased intersegmental fasciculates, branches and commissures (**Figure 3B-D**). This phenotype suggests that loss of dWdr45 affects the regulation of axon guidance, branching, and connectivity during embryogenesis. Along the same lines, the larval NMJ exhibited an increased number of branches but an identical number of synaptic terminal formation (boutons), a phenotype observed in both heterozygous and homozygous *dwdr45 KO flies* when compared to control flies (**Figure 3 E-G**). The specific increase in NMJ branches without a change in bouton number suggests that while axonal arborization is enhanced or dysregulated, synaptic terminal formation remains tightly controlled. Overall, these structural abnormalities at both embryonic and larval stages reinforce the role of dWdr45 in regulating neuronal morphogenesis and structural integrity, which are critical for proper circuit formation and motor function.

**Figure 3.**
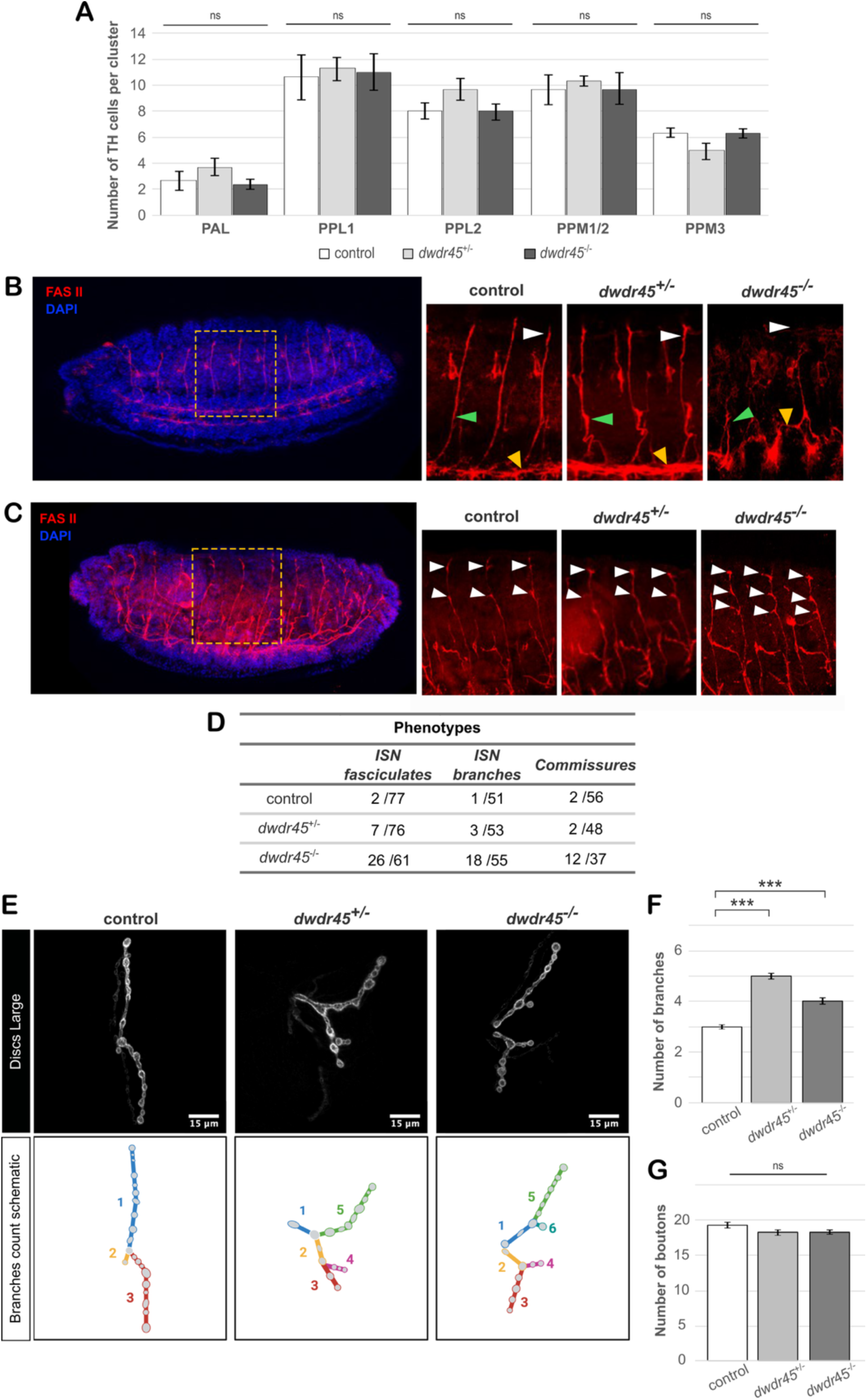
Neuronal morphological defects in the embryonic nervous system and larval neuromuscular junction of *dwdr45* KO flies. (**A**). Quantification of tyrosine hydroxylase (TH) positive cells for dopaminergic neurons clusters was realized by immunofluorescence of TH in brains of 8-day-old adult control, heterozygous *dwdr45* KO or homozygous *dwdr45* KO flies. Statistical analyses were carried out in 3 independent experiments (n=3) with ANOVA1: ^ns^no significant differences. (**B** and **C**) Immunofluorescence of FAS II in control, heterozygous *dwdr45* KO or homozygous *dwdr45* KO embryos reveals (**B**) structure of the intersegmental nerve (ISN, white arrowheads), segmental nerve (SN, green arrowheads), and commissure roots (yellow arrowheads), as well as in (**C**) fasciculations (white arrowheads). (B) and (C) are embryos captured from different focal plans. (**D**) Number of embryos showing alteration of these structures. (**E**) Immunofluorescence of discs large reveals postsynaptic boutons of L3 larval NMJ structure in muscle 12 from A2-A4 segments of control, heterozygous *dwdr45* KO or homozygous *dwdr45* KO flies with its schematic version to highlight the branches count (F), and number of postsynaptic boutons (G). Statistical analyses were carried out in 3 independent experiments (n=3) with ANOVA1: ns, no significant difference, and ***p-value<0.001.

### Fertility and lifespan reduction in *dwdr45* KO flies

In addition to the impaired locomotion and seizure-like behavior, we found that homozygous *dwdr45* KO are weak fly stocks, potentially resulting from both reduced fertility and lifespan. To assess fertility, we quantified the number of eggs laid that successfully developed into larvae, alongside counts of unfertilized (white) and non-viable (dark) eggs (**Figure 4 A, B**). Homozygous *dwdr45* KO flies showed a marked reduction in both the total number of eggs laid and hatching rates compared to controls. Moreover, the proportion of unfertilized eggs was increased in both heterozygous and homozygous *dwdr45* KO compared to control flies. Notably, *dwdr45* KO females laid over 40% of non-viable eggs. These findings suggest a fertility impairment and embryonic lethality induced by homozygous loss of *dwdr45*. To assess lifespan, we conducted longevity assays. Heterozygous *dwdr45* KO flies exhibited a nearly 50% reduction in the mean adult lifespan relative to that of control flies, while homozygous *dwdr45* KO flies exhibited a severe reduction of lifespan with a mean lifespan of ≈10 days compared to ≈50 days in control flies (**Figure 4C, D, E**).

**Figure 4.**
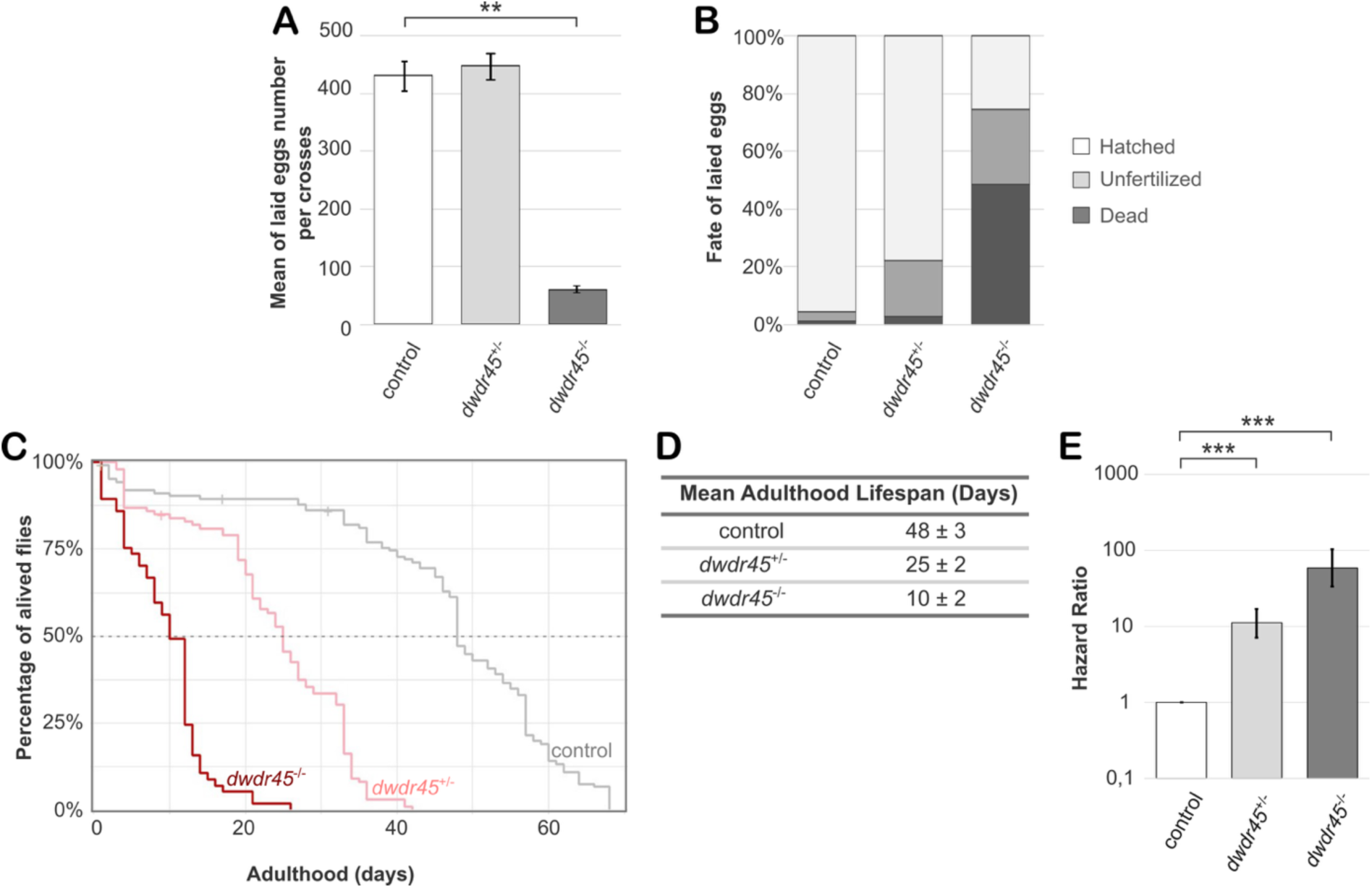
Fertility defects and reduced lifespan in *dwdr45* KO flies. (**A** and **B**) Fertility was assayed by (**A**) quantifying the number of eggs laid and (**B**) their fate unfertilized, dead and subsequent larval hatching. Control, heterozygous *dwdr45* KO or homozygous *dwdr45* KO female flies were crossed with control male flies. Statistical analyses were carried out in three independent experiments (n=3) with ANOVA1/post-hoc: **p-value<0.01. (**C**, **D** and **E**) Lifespan assays were realized in control, heterozygous *dwdr45* KO or homozygous *dwdr45* KO female flies with Kaplan-Meier analysis. (**C**) A representative lifespan experiment out of three independent experiments (n=3) is shown. (**D**) The mean adulthood lifespan and (**E**) the hazard ratio (probability of fly death event) are shown in the indicated genetic backgrounds. Statistical analyses were carried out with Coxph test: ***p-value<0.001.

### Iron dyshomeostasis in *dwdr45* KO flies

Iron dyshomeostasis has been reported in fibroblasts derived from BPAN patient tissues and in HeLa cells overexpressing mutant form of *wdr45* (31, 32). In contrast, most *wdr45* KO mouse models do not exhibit significant iron dysregulation (13, 16, 17). To investigate iron homeostasis, we measured ferritin protein levels and total iron content in control and *dwdr45* KO flies. Ferritin status was assessed by quantifying the protein levels of Fer1HCH and Fer2LCH by western blotting (**Figure 5A**-**D**). We observed elevated levels of both Fer1HCH and Fer2LCH proteins in homozygous *dwdr45* KO flies relative to controls, with Fer1HCH increase detected in newly eclosed (1-day-old) flies and Fer2LCH increase detected in 8-day-old adults.

**Figure 5.**
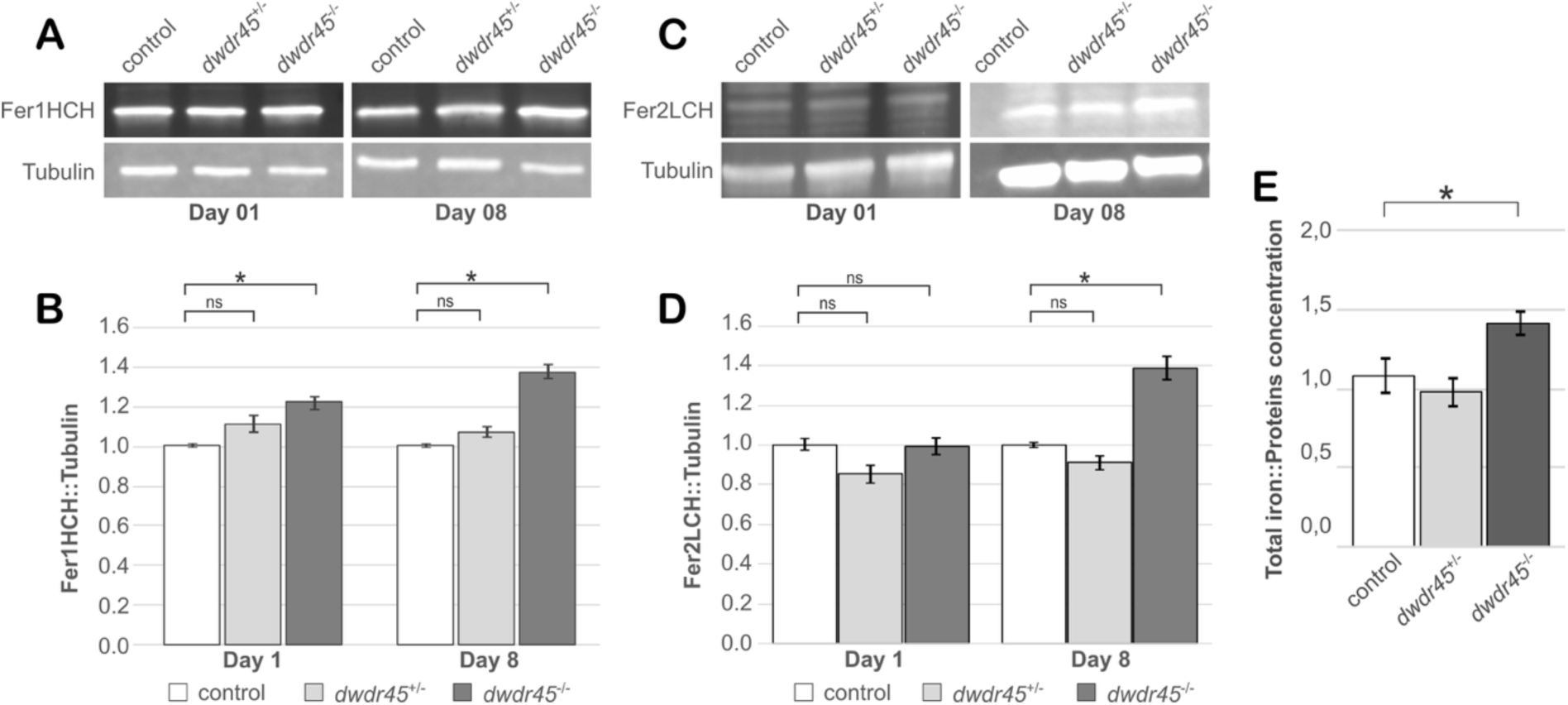
Loss of *dwdr45* alters ferritins expression and disrupts iron balance in *dwdr45* KO flies. (**A**) Fer1HCH proteins are detected by Western Blotting in 1- and 8-day-old adult control, heterozygous *dwdr45* KO or homozygous *dwdr45* KO flies. (**B**) The ratio of Fer1HCH relative to tubulin protein levels is normalized to that of control flies. Statistical analyses were carried out in four independent experiments (n=4) with ANOVA2/post-hoc test: ns: no-significant, *p-value<0.05. (**C**) Fer2LCH proteins are detected by western blotting in 1- and 8-day-old adult control, heterozygous *dwdr45* KO or homozygous *dwdr45* KO flies. (**D**) The ratio of Fer2LCH relative to tubulin protein levels is normalized to that of control flies. Statistical analyses were carried out in four independent experiments (n=4) with ANOVA2/post-hoc test: ns: no-significant, *p-value<0.05. (**C**) Ratio of iron on protein concentration was quantified by a ferrozine assay in 1-day-old adult control, heterozygous *dwdr45* KO or homozygous *dwdr45* KO flies. Statistical analyses were carried out in three independent experiments (n=3) with ANOVA1/post-hoc test: *p-value<0.05.

In *Drosophila*, it has been proposed that ferritin levels increase following iron feeding in the gut of third instar larvae (33). Consistently, we showed that Fer1HCH and Fer2LCH levels increased in adult *Drosophila* when fed with an iron rich diet (**Figure S4**). We thus hypothesized that the increase in ferritin protein levels in homozygous *dwdr45* KO flies may be due to iron accumulation, a characteristic of BPAN (15). To test this, we conducted a ferrozine assay to quantify the total iron content in *dwdr45* KO and control flies (**Figure 5E**). The results indicated a significant increase in total iron content in 1-day-old homozygous *dwdr45* KO flies compared to controls, supporting the notion that dWdr45 is essential for maintaining iron homeostasis in *Drosophila*. Together, these findings demonstrate that homozygous *dwdr45* KO flies exhibit iron dyshomeostasis, evidenced by elevated levels of Fer1HCH, Fer2LCH, and total iron.

### Autophagy dysregulation in *dwdr45* KO flies

Autophagy defects have been documented in human cellular and animal models such as mouse and *Dictyostelium* models with mutations in *WDR45* or its homologs, respectively (2, 34). During the autophagy process, the accumulation of autophagic vacuoles can be detected with staining of ATG8a in flies or LC3 in mammals (35). To assess autophagy levels, we first measured ATG8a protein accumulation by western blot using an anti-GABARAP in *Drosophila* head extracts (**Figure 6A, B**). An increase in the lipidated ATG8a-II form is commonly used as an indicator of autophagic vacuole build-up. We observed an increased ATG8a-II/ATG8a-I ratio in homozygous *dwdr45* KO compared to control flies, indicating a higher level of autophagic vacuoles. To further evaluate autophagy activity, we utilized the *Drosophila* retina as a model system to study autophagy in the nervous system, as previously described (24, 36). The pan-retinal knockdown of *dwdr45* by RNAi resulted in the accumulation of GFP dots in flies expressing a transgenic GFP::LC3 construct in photoreceptor cells (**Fig. S5A, B**). The quantification of GFP::LC3 dots in the retina showed that the accumulation in *dwdr45* knockdown flies was comparable to that seen with the overexpression of the Atro75QN mutant that causes autophagic vacuole accumulation due to a blockage in autophagic flux in the *Drosophila* retina (36, 37).

**Figure 6.**
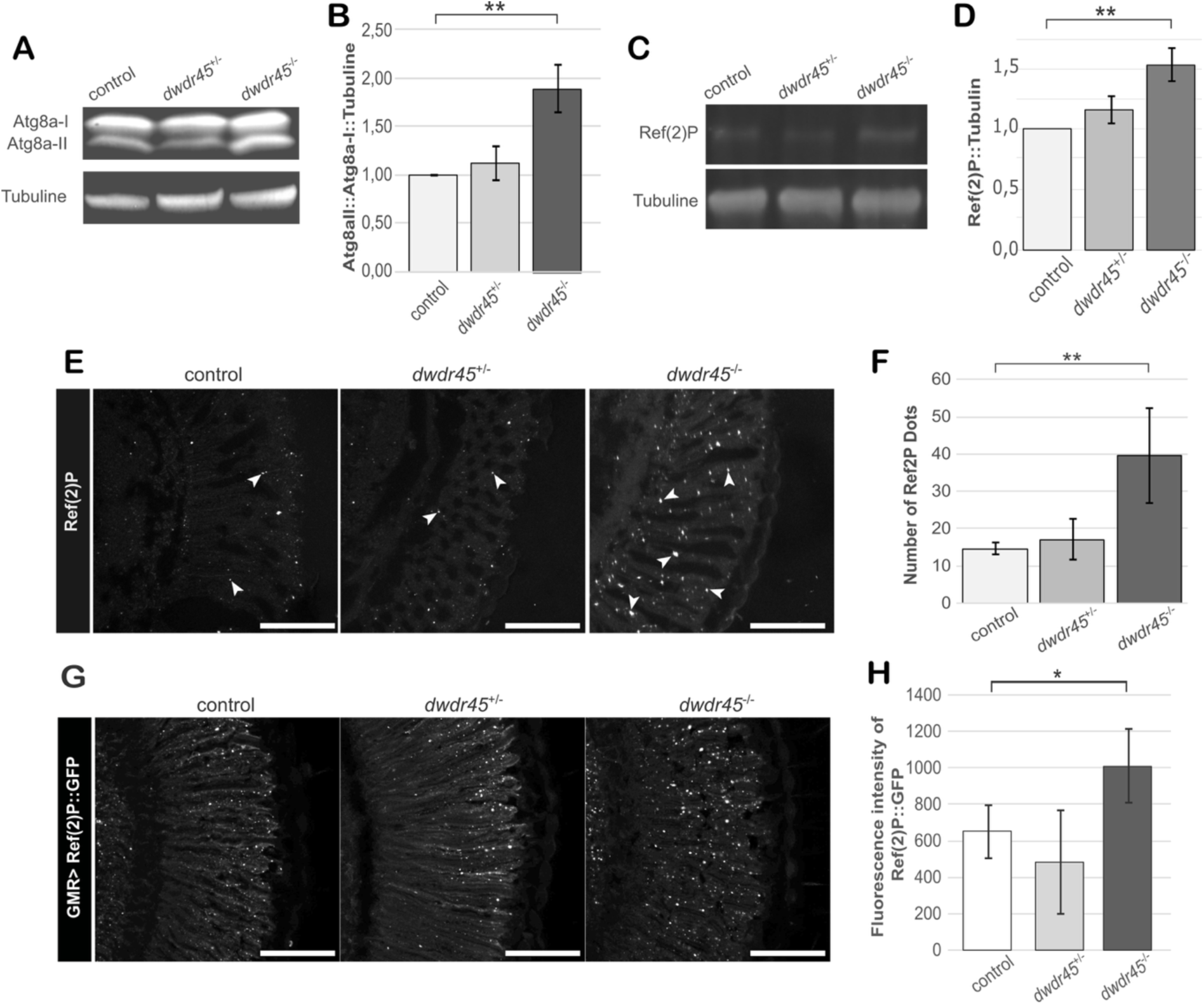
Loss of *dwdr45* deficiency impairs autophagic flux in adult flies. (**A**) Atg8a-II/Atg8a-I proteins are detected by Western Blotting in 8-day-old adult control, heterozygous *dwdr45* KO or homozygous *dwdr45* KO flies. (**B**) The ratio of Atg8a-II/Atg8a-I relative to tubulin protein levels is normalized to that of control flies. Statistical analyses were carried out in three independent experiments (n=3) with ANOVA1/post-hoc test: **p-value<0.01. (**C**) Ref(2)P protein is detected by Western Blotting in 8-day-old adult control, heterozygous *dwdr45* KO or homozygous *dwdr45* KO flies. (**D**) Ref(2)P relative to tubulin protein levels is normalized to that of control flies. Statistical analyses were carried out in three independent experiments (n=3) with ANOVA1/post-hoc test: *p-value<0.05. (**E**) Immunolabelling of Ref(2)P protein in the retina of 15-day-old adult control, heterozygous *dwdr45* KO or homozygous *dwdr45* KO flies. (**F**) The number of Ref(2)P dots relative to retina surface area was quantified. Statistical analyses were carried out in three independent experiments (n=3) with ANOVA1/post-hoc test: **p-value<0.01. Scale bars represent 50 μm. (**G**) Immunolabelling of GFP adult retina overexpressing Ref(2)P::GFP under the pan-retinal GMR driver in 15-day-old control (*;GMR-Gal4;UAS-Ref(2)P::GFP*), heterozygous *dwdr45* KO or homozygous *dwdr45* KO flies (*;GMR-Gal4;UAS-Ref(2)P::GFP,dwdr45^KO^*). (**H**) Quantification of GFP fluorescence. Statistical analyses were carried out in three independent experiments (n=3) with ANOVA1/post-hoc test: *p-value<0.05. Scale bars represent 50 μm.

We then assessed autophagic flux, which reflects the efficiency of autophagy in degrading cellular components. For this purpose, we examined Ref(2)P/p62, a protein that is typically degraded in autolysosomes during autophagy (36, 38). We observed an accumulation of Ref(2)P/p62 protein by Western blot from *Drosophila* head extracts (**Figure 6C, D**) and by immunofluorescence in *Drosophila* retina (**Figure 6E, F**). Additionally, increased Ref(2)P was detected in retina expressing *dwdr45* RNAi, comparable to the Ref(2)P accumulation seen in retina overexpressing *Atro75QN* (**Fig. S5C, D**).

We further checked whether Ref(2)P accumulation in homozygous *dwdr45* KO retina results from impaired autophagic degradation. For this, we overexpressed Ref(2)P::GFP specifically in photoreceptors using the pan-retinal GMR driver (**Figure 6G, H**). Ref(2)P::GFP immunostaining was advantageous over endogenous Ref(2)P as it is unaffected by ROS-driven *Ref*(*2*)*P* promoter regulation (36, 39), and serves as a tool to probe the autophagy dysfunction. Ref(2)P::GFP levels were significantly elevated in homozygous *dwdr45* KO retina compared to that of controls, indicating that Ref(2)P accumulation is indeed due to defective autophagy. Together, the accumulation of both ATG8a and Ref(2)P/p62 strongly suggest an impairment of autophagic flux in *dwdr45* KO flies.

### Neuronal induction of autophagy improves seizure-like behavior and longevity but not locomotion deficits in dwdr45 KO flies

Autophagy is a central mechanism to ensure neuronal health and lifespan in *Drosophila* (12, 40, 41). To determine whether the observed deficits in autophagy in *dwdr45* KO flies contribute to their locomotion and lifespan phenotypes, we performed neuronal autophagy induction experiments by overexpressing Atg1 or human Lamp2A, as previously used to induce autophagy in *Drosophila* (36, 42, 43). We first observed that the pan-neuronal overexpression of Atg1 or Lamp2A could decrease the Atga8II/Atg8aI ratio in *dwdr45 KO* flies to a level similar to that of control flies (**Figure 7A, B and C, D**) suggesting that these genetic manipulation provoke autophagic induction. However, neither Atg1 overexpression nor Lamp2A expression affected the locomotion of *dwdr45* KO flies in 1-, 4-, 8- and 15-day-old flies (**Figure 8 A, B**). This result suggests that dWdr45 may possess at least one additional function, separate from autophagy, that is essential for normal locomotion. In contrast, the overexpression of Atg1 or Lamp2A was sufficient to rescue the seizure-like behavior exhibited by *dwdr45* KO flies (**Figure 8 C-J**). Interestingly, while Atg1 and Lamp2A overexpression effectively restored seizure-like phenotypes in both heterozygous and homozygous *dwdr45* mutants, their overexpression in wild-type flies actually exacerbated the seizure-like behavior (**Figure 8 C, D**). This could indicate that although autophagy deficiency underlies the seizure-like phenotype in *dwdr45 KO* flies, maintaining a precise autophagy balance is critical, as excessive induction of autophagy can provoke seizures in control flies. Finally, we showed that increased autophagy mediated by overexpression of Atg1 or expression of human Lamp2A improved the lifespan of *dwdr45* KO flies (**Figure 8E-J**), which indicates that neuronal induction of autophagy is beneficial in *dwdr45* KO flies. As shown previously, Atg1 overexpression provoked lifespan extension in control flies (42). Surprisingly, in this experiment, control flies had a shorter lifespan than *dwdr45* KO heterozygous flies, which was attributed to a background dominant mutation carried on the E*lav-Gal4;;* line, possibly due to a floating balancer (data not shown). Nevertheless, we observed that Atg1 or Lamp2A overexpression was more beneficial in homozygous compared to heterozygous *dwdr45* KO, which suggests that the reduced longevity in *dwdr45* KO flies is, at least in part, due to autophagy defects. Collectively, the neuronal induction of autophagy in *dwdr45* mutants differentially influences locomotion, seizure-like behavior, and longevity, highlighting distinct roles of autophagy in the various *dwdr45* KO phenotypes and suggesting the existence of autophagy-independent functions of dWdr45 in the maintenance and function of the *Drosophila* nervous system.

**Figure 7.**
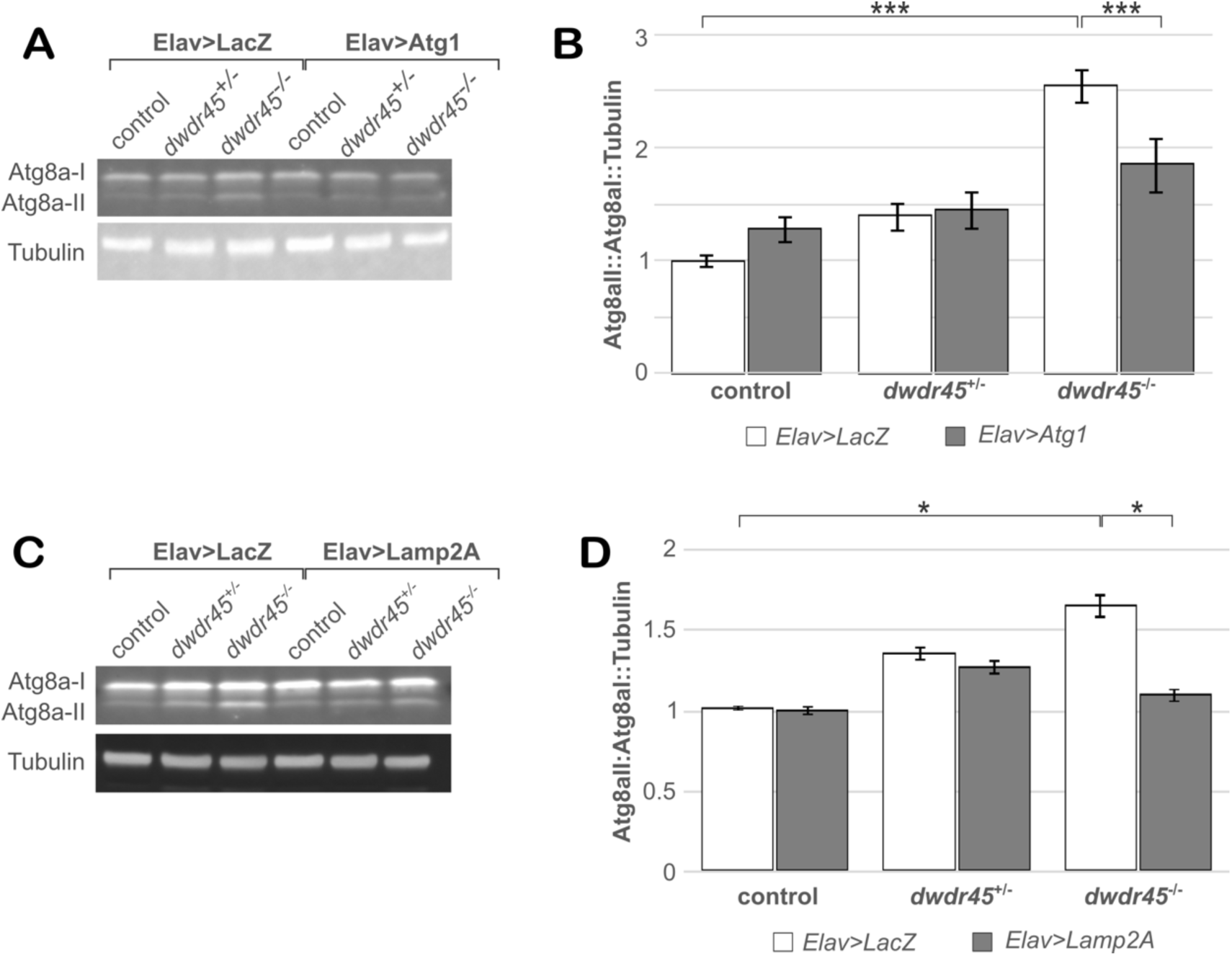
Atg1 or Lamp2A restore autophagy defects in *dwdr45* KO flies. (**A**) Atg8a-II/Atg8a-I proteins are detected by Western Blotting in 8-day-old adult control (*Elav-Gal4;;*), heterozygous *dwdr45* KO or homozygous *dwdr45* KO flies (*Elav-Gal4;;dwdr45^KO^*) expressing *LacZ* or *Atg1* under *Elav* driver. (**B**) The ratio of Atg8a-II/Atg8a-I relative to tubulin protein levels is normalized to that of *Elav>LacZ* flies. Statistical analyses were carried out in three independent experiments (n=3) with ANOVA1/post-hoc test: **p-value<0.01, ***p-value<0,001. (**C**) Atg8a-II/Atg8a-I proteins are detected by Western Blotting in 8-day-old adult control (*Elav-Gal4;;*), heterozygous *dwdr45* KO or homozygous *dwdr45* KO flies (*Elav-Gal4;;dwdr45^KO^*) expressing *LacZ* or human *Lamp2A* under *Elav* driver. (**D**) The ratio of Atg8a-II/Atg8a-I relative to tubulin protein levels is normalized to that of *Elav>LacZ* flies. Statistical analyses were carried out in three independent experiments (n=3) with ANOVA1/post-hoc test: *p-value<0.01.

**Figure 8.**
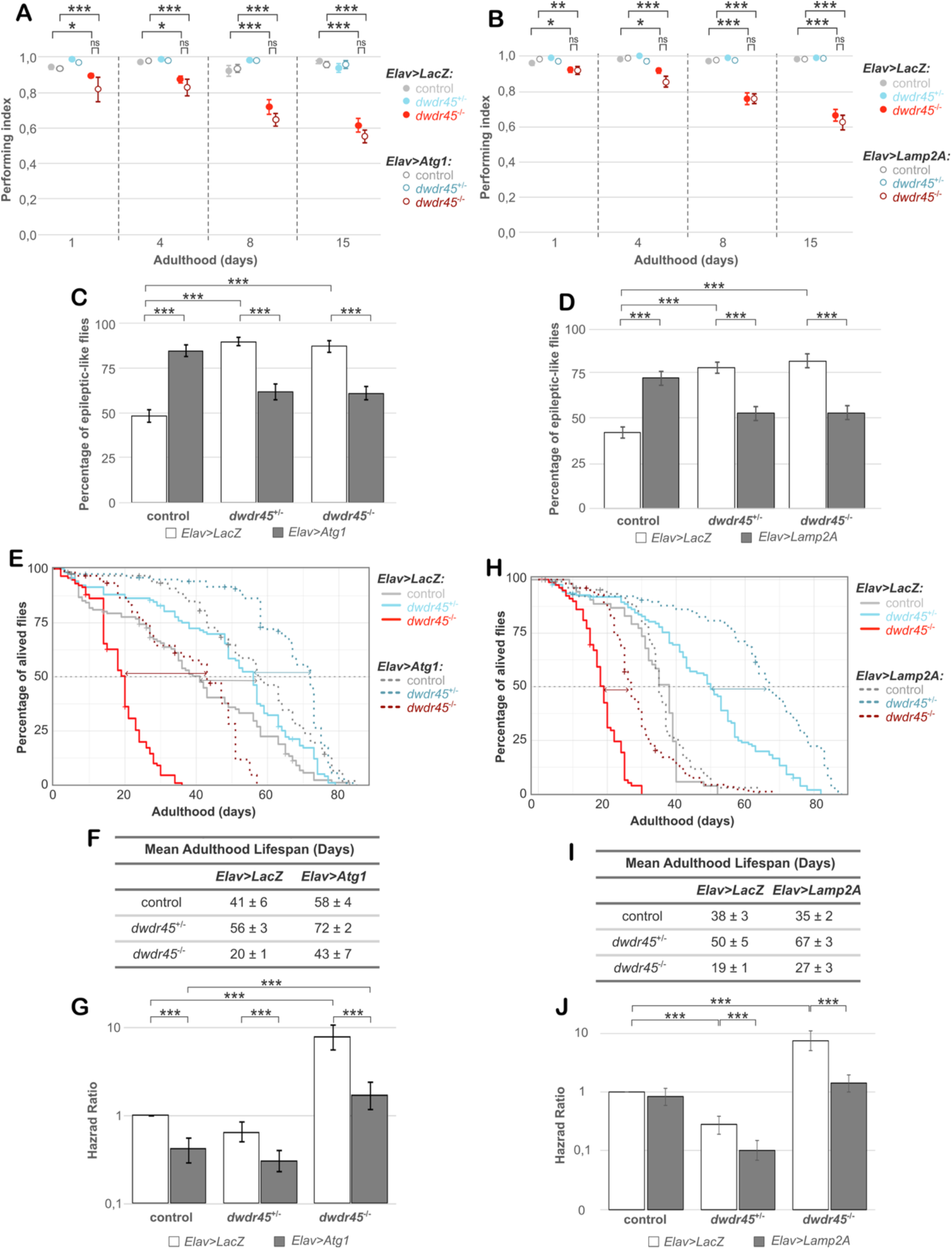
Neuronal overexpression of *Atg1* or *Lamp2A* improves seizure-like phenotype and lifespan but not locomotion in *dwdr45* KO flies. Neuronal overexpression of *LacZ* or *Atg1* or *Lamp2A*, under the pan-neuronal driver *Elav*, in control (*Elav-Gal4;;*), heterozygous *dwdr45* KO or homozygous *dwdr45* KO flies (*Elav-Gal4;;dwdr45^KO^*). (**A**) Locomotion SING assays were realized in 1-, 4-, 8- and 15-day-old adult flies overexpressing *LacZ* or *Atg1*. Statistical analyses were carried out on performing index in three independent experiments (n=3) with ANOVA2/post-hoc test: ns, no significant, *p- value<0,05, ***p-value<0,001. (**B**) Locomotion SING assays were realized in 1-, 4-, 8- and 15- day-old adult flies overexpressing *LacZ* or *Lamp2A*. Statistical analyses were carried out on performing index in three independent experiments (n=3) with ANOVA2/post-hoc test: ns, no significant, *p-value<0,05, **p-value<0,01, ***p-value<0,001. (**C, D**) Percentage of flies exhibiting seizure-like phenotype, described as loss of posture, leg shaking, random wing flapping, spinning and uncontrolled flight, was measured in 4-day-old flies overexpressing *LacZ*, or *Atg1* (**C**) or *Lamp2A* (**D**) under the pan-neuronal driver *Elav*. Statistical analyses were carried out in 3 independent experiments (n=3) with ANOVA1: ***p-value<0.001. (**E**, **F** and **G**) Lifespan experiments were realized in adult flies overexpressing *LacZ* or *Atg1*. (**E**) Representative analysis out of three independent experiments is shown (n=3). (**F**) Associated mean adulthood lifespan and (**G**) hazard ratio (probability of fly death event). Statistical analyses carried out with Coxph test (***p-value<0.001). (**H**, **I** and **J**) Lifespan experiments were realized in adult flies overexpressing LacZ or Lamp2A. (**H**) Representative analysis out of three independent experiments is shown (n=3). (**I**) Associated mean adulthood lifespan and (**J**) hazard ratio (probability of fly death event). Statistical analyses carried out with Coxph test (***p-value<0.001).

## Discussion

In this study, we establish a novel *Drosophila* model to study BPAN. We demonstrate that homozygous *dwdr45* KO flies display a range of cellular and behavioural phenotypes that phenocopy those observed in BPAN animal models and disease. Specifically, *dwdr45* KO flies exhibit autophagy dysfunction and iron accumulation at the cellular level, which correlate with locomotor impairment, seizure-like behavior and reduced lifespan at the organismal level. As today, seven mouse BPAN models have been described in the literature (13, 16, 17, 28, 44–46). All of them, except Noda et al., show at least autophagic defects and locomotor dysfunction, sometimes with more cellular (mitochondrial, ER stress) and organismal (memory, social ability, neurodegeneration) characterizations. Notably, only one mouse model exhibits iron accumulation and seizure-induced phenotype (13), a finding consistent with the *dwdr45*KO fly model we have established. Importantly, Noda et al. 2021, remains the only study specifically investigating the neurodevelopmental defects in a BPAN mouse model (46). They reported synaptogenesis abnormalities with arborization defects, which also seems to be impaired in *Drosophila* BPAN model. Overall, *dwdr45 KO* fly model recapitulates key phenotypes described across mouse models and consolidates them within a single organism. The *Drosophila* BPAN model appears as a valuable system to investigate the causal relationships between the various cellular defects and the resulting organismal phenotypes and serves as a powerful tool for pharmacological testing prior to mammal studies.

Remarkably, pan-neuronal overexpression of Atg1 or human Lamp2A, which induces autophagy, were able to improve the seizure-like behavior and to extend lifespan in *dwdr45* KO flies. Mammalian WDR45 participates in phagophore elongation, autophagosome maturation, and the autolysosome maturation step (2). Under nutrient-rich conditions, the WDR45 complex associates with ATG2, AMPK, and ULK1/ATG1. Upon starvation, the WDR45/WIPI4–ATG2 complex dissociates and relocates to nascent autophagosomes (6), indicating that WDR45 collaborates with ATG1 during the initiation phase of autophagy. Therefore, dWdr45 may likewise act with Atg2 and Atg1 at the onset of autophagy, consistent with the presence of a conserved Atg2-binding motif in its protein sequence (**Figure 1**). Such a function could account for the rescue of *dwdr45* KO phenotypes by Atg1 overexpression. Alternatively, the positive effects of Atg1 overexpression in *dwdr45* KO flies may result from a partial impairment of autophagy upon *dwdr45* loss, permitting Atg1-driven enhancement of the autophagic process and improved cellular adaptation.

The rescue of the longevity and the seizure-like phenotypes mediated by Lamp2A expression in *dwdr45* KO flies could be explained by its function on macroautophagy or endosomal microphagy. Indeed, while Lamp2A is required for chaperone mediated autophagy (CMA) in mammalian cells (47), it is involved in ESCRT-dependent endosomal microautophagy in *Drosophila* (43). *Drosophila* lacks CMA but instead uses endosomal microphagy as a selective autophagic process (48). Furthermore, it was shown that human Lamp2A induces macroautophagy by promoting Atg5 expression (43). In our experiments, we demonstrated that Lamp2A expression rescues the increased Atg8a-II/Atg8a-I ratio to normal levels in *dwdr45* KO flies, indicating induction of macroautophagy. This suggests that Lamp2A-mediated neuronal induction of macroautophagy, similar to Atg1, rescues the longevity and the seizure-like phenotypes. Whether the endosomal microautophagy also contributes to the rescue of aging and seizure-like phenotypes observed in *dwdr45* KO remains to be elucidated. Although autophagy and aging are well studied, and impaired autophagy is well known to contribute to aging (49), the role of autophagy in epilepsy remains poorly defined despite emerging evidence. Importantly, given that epilepsy in BPAN is frequently resistant to pharmacological treatment, the *dwdr*45 KO fly model represents a valuable system for identifying new anti-epileptic compounds.

We found that the neuronal induction of autophagy via overexpression of Atg1 or expression of Lamp2A rescued the seizure-like phenotype and reduced lifespan but failed to improve locomotor deficits in *dwdr45* KO flies. Furthermore, we showed that neuronal overexpression of *dwdr45* is sufficient to restore the locomotion phenotype, whereas neuronal overexpression of Atg1 does not rescue the locomotion defects in *dwdr45* KO flies. Given that the locomotor defects of *wdr45*KO flies have a strong neuronal component, if the effect of d*wdr45* loss were directly linked to its role in neuronal autophagy, a rescue would be expected. However, it remains possible that the level of autophagy induction is insufficient or that other cell types contribute to the phenotype. Our results also suggest that *dwdr45* may also have autophagy-independent functions, which are required for locomotion. This is supported in two recent studies, proposing that autophagy-independent roles of WDR45 could lead to additional cellular dysfunctions. The first study showed that WDR45 loss triggers ATG2 recruitment to mitochondria, inducing ferroptosis *via* an autophagy-independent pathway (8). The second study identified a specific role of WDR45, via its WD5 domain, in stress granule disassembly, which correlates with early disease onset (50). The partial rescue of phenotypes by overexpression Atg1 or expression of Lamp2A highlights the therapeutic relevance of targeting autophagy in BPAN and also suggests the need for complementary therapies targeting non-autophagic roles of dWdr45.

The lack of neurodegeneration of dopaminergic neurons in the *dwdr45* KO fly model could suggest that other neuronal subpopulations are affected or that more subtle structural defects of neurons are present in the adult with age or developing nervous system. The latter is supported by our observation that neurodevelopmental defects, such as ectopic neuronal branches and fasciculations are observed in both the embryonic nervous system and larval NMJ. A thorough characterization of the arborization of the different neuronal subpopulations will be necessary to further characterize the neuronal and autophagy functions of dWdr45 in locomotion, seizure and aging.

Iron dysregulation remains a topic of debate in BPAN cellular and animal models. While several cellular models report iron accumulation in *wdr45*-deficient cells, measurements of ferritin levels and ferritinophagy vary across cell types (3, 31, 32, 51). Additionally, iron accumulation is not consistently detected in *wdr45* KO mice (13, 16, 17), despite being a key feature of BPAN disease. These findings suggest that iron dysregulation may occur in BPAN cellular models, but its physiological relevance in whole organisms is still unclear. We found that BPAN flies accumulate iron, which correlates with elevated levels of Fer1HCH and Fer2LCH. Although insect ferritins are secreted, unlike their mammalian counterparts, their abundance reliably reflects intracellular iron content. Consistently, previous findings have shown that *Drosophila* ferritin levels increase in the gut of third instar larvae fed an iron-enriched diet (33). Furthermore, we detected elevated Fer1HCH and Fer2LCH levels in adult flies under similar rich iron-diet conditions. However, the comparison of the detailed mechanisms regulating iron homeostasis in *wdr45* KO between flies and mammalian could be limited by the different transport iron mechanisms and the fact that ferritins are secreted in flies. Nonetheless, *Drosophila* as mammalian ferritins, play a key role in iron detoxification by storing excess labile iron and preventing tissue damage (20, 52, 53). *Drosophila* thus provides a valuable model for investigating how iron accumulation contributes to the cellular and organismal defects associated with *dwdr45* KO, which warrants further study. Furthermore, it is still unknown whether ferritinophagy could play a role in the regulation of ferritin protein levels in *Drosophila* but a previous study reported that ferritin levels are largely independent of autophagy in *Drosophila* (54). Whether dWDR45 regulates iron homeostasis in an autophagy independent manner remains to be investigated.

Lastly, dWdr45 is the only homolog for both WDR45 and WDR45B in mammals. The ability of human h*WDR45* to rescue the locomotion defect in *dwdr45* KO flies supports *dwdr45* KO flies as a valuable model for studying conserved functions of WDR45 in BPAN. Further research could explore the role of human WDR45B in flies, which would be highly relevant to studying neurodevelopmental pathologies associated with WDR45B mutations. Such conditions, including developmental delay, microcephaly, seizures, and neurological deterioration, partially overlap with BPAN disease phenotypes (55).

## Materials and Methods

### *Drosophila* culture and strains

Flies were grown on regular cornmeal-yeast medium at 25°C with a 12 h light/dark cycle. The fly stocks were obtained as followed: *CG11975^KO-kG4^* (BL97305), *UAS-GFP (on chromosome III)* (56), *Elav-Gal4 (on X)*, *UAS-WDR45.HA* (on II) (BL84770), *UAS-αSynuclein* (on III) (BL8146), *sGMR-Gal4 (on II)*, *UAS-LacZ* (on III) (BL1777) and *UAS-Atg1* (on II) (BL51654) were from Bloomington Drosophila Stock Center. *UAS-LacZ* RNAi (on II) (51446) and *UAS-dwdr45* RNAi (on III) (GD26799) were from Vienna Drosophila Resource Center. *UAS-GFP::dW*w*dr45* (on III) was provided by P. Verstreken, *UAS-Atro75QN (on II)* was provided by M. Fanto (KCL, UK), Para^BSS1^ (on X) was provided by R. Baines (University of Manchester, UK), *UAS-GFP-LC3* (on II) was provided by H. Stenmark (University of Oslo, NO), *UAS-Ref*(*2*)*P::GFP* (on III) was provided by T. Neufeld (University of Minnesota, USA) and *UAS-Lamp2A* (on II) (43).

Additionally, several strains were crossed with the *wdr45* KO line to perform experiments in a *dwdr45* mutant context. Complete genotypes are provided in **Table S1**.

Except when specified in the figure legends, control flies correspond to the genotype *(w^1118^;;)*; heterozygous flies to *(w^1118^;; dwdr45^KO^/TM6B, Tb)*; and homozygous flies to *(w^1118^;; dwdr45^KO^)*.

### Sequences identification and analysis

The *Drosophila* homolog of WDR45/WIPI4 was identified by performing a protein BLAST search on NCBI (https://blast.ncbi.nlm.nih.gov). To perform phylogenetic analysis, we used amino acid sequences of paralogs and orthologs of WDR45/WIPI4 in *Homo sapiens*, *Mus musculus*, *Drosophila melanogaster*, *Dictyostelium discoideum* and *Saccharomyces cerevisiae* to analyze multiple sequence alignment using ClustalX with SeaView software. To define the evolutionary relationship, the phylogenetic tree was constructed using the PhyML method on SeaView software. A schematic representation of the 3D structures colorized depending on sequence conservation was created using ChimeraW software with 3D structures of WDR45 and dWdr45 downloaded from Swiss-Model, and sequence alignment previously performed in SeaView software.

### Generation of *dwdr45* KO flies and genotyping

To generate whole body *dwdr45* KO mutant flies, we used TRIP-CRISPR lines from Bloomington Drosophila Stock Center. Firstly, flies expressing Cas9 in germline cells through nanos driver (BL54593) were crossed with flies expressing *dWdr45* guide RNA (BL83546). The expression of both guide RNA and Cas9 in first progeny (F1) males induces a deletion at position 222 of *dwdr45* gene in germline cells. To obtain whole body *dwdr45*^+/-^ mutant flies, F1 males were then crossed with flies carrying *TM6B,Tb* (*w^1118^;;TM6B,Tb/Sb*)balancer 50 flies were collected from this second progeny (F2) which were individually crossed with balanced flies to eliminate Cas9 and guide RNA transgene.

To confirm *dwdr45* deletion, genotyping was performed before each crosses. After genomic DNA extraction, *dWdr45* gene was amplified by touchdown PCR using the following primers: forward 5’-CATATGGAAACGGGCTGCTC-3’, reverse 5’-ATCCCACACGATCACCTTG-3’. Since *dwdr45* guide RNA targets the digestion site of BsaWI enzyme, DNA with Cas9-mediated deletion lacks this restriction site. Once *dwdr45* mutation was confirmed by restriction, *dWdr45* gene was sequenced to identify the precise deletion. We selected flies carrying a deletion of 7 nucleotides from nucleotide 222 to 229. This deletion induces a frameshift and a premature stop codon leading to a protein of 38 amino acids. This protein is predicted to be non functional due to the loss of the seven-beta blade-propeller structure and the loss of interaction domain with autophagy key regulators. Finally, to cross out guide RNA and Cas9 transgenic constructs, flies were crossed multiple times with flies carrying TM6B,Tb balancer (*w1118;;dwdr45KO/TM6B,Tb*).

### Adult brain immunostaining

Brains from 8-day-old adult female flies were dissected in Schneider medium (Thermo Fisher Scientific, 21720001) and stored in Schneider + 1% paraformaldehyde (PFA [Euromedex, EM-15710]) until end of dissections. Brains were fixed in Schneider + 4% PFA for 25 min at room temperature followed by three-times 20 min washes in washing solution (PBS1X [Thermo Fisher Scientific, 14200083] + 0.3% Triton-X-100 [Sigma-Aldrich, T8787] + 5 mg/mL BSA [Euromedex, 04-100-812-E]). Saturation was realized over 1 h in blocking solution (washing solution + 4% NGS [Sigma-Aldrich, G9023]). Brains were incubated in primary antibodies overnight at 4°C with rabbit anti-GFP (1:200 [Thermo Fisher Scientific, A6455]), mouse anti-Elav (1:400 [Developmental Studies Hybridoma Bank, 7E8A10]) or rabbit anti-tyrosine-hydroxylase (1:500 [Sigma-Aldrich, AB152]) diluted in blocking solution. Brains were washed three times for 20 min each before secondary antibodies incubation for 3 h at room temperature with anti-Rabbit-A488 (1:400 [Thermo Fisher Scientific, A-11008]) or anti-Mouse-A546 (1:400 [Thermo Fisher Scientific, A-11030]) diluted in blocking solution. Finally, brains washed three times for 20 min each and mounted in Vectashield (Eurobio Scientific, H-1000) on bridge slides. Confocal microscopy was performed using a Zeiss LSM800 microscope.

Quantification of dopaminergic neurons was performed manually by counting TH-positive cells within each cluster. For each brain, the mean value of the counts from the right and left cortex was calculated.

### SING Assay

Locomotion ability was measured using SING assay. Females were preferred as they perform more robustly the locomotion assay. For each condition, 6 groups of 18 adult females were harvested within one day of adult hatching. 30 min before the SING assay, flies were left at room temperature for habituation. Each group was then placed in a column (25 cm long and 1 cm in diameter). First, flies underwent three training sessions in which they were gently tapped down and let climb to the top of the column. Second, flies were assayed by gently tapping them down and counting the number of flies reaching the top of the column (n_up_ [above 18 cm]) and those staying at the bottom end (n_down_ [below 2.5 cm]) after 30 sec. Each group was assayed 3 times with 2 min intervals. The mean of the 3-times assays was used to calculate a performing index: ½[(n_tot_+n_up_-n_down_)/n_tot_] where n_tot_ is the total number of flies. For each condition, the result is the mean ±SEM of the 6 groups.

### Bang-sensitive assay

Seizure-like phenotype was assessed using mechanical stimulation, also referred to as bang-sensitive assay. 4-day-old post-eclosion female flies were collected under CO2 and distributed into vials containing five flies each, one day prior to the experiment, allowing recovery from anaesthesia. Five minutes before testing, the flies were gently moved to empty vials. To induce seizure-like phenotype, the vials were placed on a standard laboratory vortexer set to maximum speed for 15s, followed by 30s of observation during recovery. Seizure manifestations such as loss of posture, leg shaking, random wing flapping, spinning and uncontrolled flight (57), were visually assessed in each fly, and the percentage of flies displaying seizure-like phenotypes was calculated for each condition. Each vial was tested three times with five flies per vial, and five vials per condition in each experiment. All experiments were performed blind to genotype and by the same investigator to ensure intra-experimental consistency. The experiments have been conducted between 8:30 and 11:30 a.m. to minimize circadian impact. For each genotype, a minimum of 75 flies from 3 independent experiments were tested.

### Embryo nervous system immunofluorescence

Embryo collection, dechorionation, devitellinization, fixation, and staining were performed as previously described (58). In brief, embryo samples were gradually rehydrated, rinsed in 1× PBS with 0.3% Triton X-100 (PBTx), and washed in PBTx (4 × 30 min). Next, they were incubated overnight at 4 °C with the following primary antibodies diluted in PBTx: mouse anti-Fasciclin II (DSHB, 1:100, 1D4) and goat anti-Horseradish Peroxidase (Jackson ImmunoResearch, 1:100, 123005021). Primary antibodies were removed by three rinses followed by four 30-minute washes in PBTx. Secondary antibodies were then applied overnight at 4 °C. The secondary antibodies used were: Alexa Fluor 555-conjugated anti-mouse (Invitrogen, 1:500) and Alexa Fluor 488-conjugated anti-goat (Invitrogen, 1:500). During secondary incubation, DAPI (Life Technologies, 1:1000) was added for nuclear staining. After staining, samples were rinsed three times, washed 4 × 30 min in PBTx, and mounted in Vectashield. Imaging was performed using a Nikon confocal laser scanning microscope, and images were processed with ImageJ/FIJI. Structural alterations, including fasculation, branches and commissures, were counted manually.

### Larval NMJs immunostaining

Third instar larvae were dissected for NMJ immunohistochemistry as previously described by Hernandez-Diaz et al. 2022 (59). Larvae were dissected in HL3 solution (110 mM NaCl, 5 mM KCl, 10 mM NaHCO3, 5 mM Hepes, 30 mM sucrose, 5 mM trehalose, and 10 mM MgCl2, pH 7.2), fixed in HL3+PFA 4% for 20 min at room temperature and permeabilized for 45 min with 0.4% Triton X-100 in PBS (PBS-T). Samples were then blocked in blocking solution (PBS-T+NGS 10%) for 30min. Immunostaining was performed with mouse anti-Dlg (1:500 [DSHB, AB-528203) overnight at 4°C followed by anti-mouse-A546 secondary antibody for 1h30, both diluted in blocking solution. Between and after antibodies incubations, larvae were washed three times for 15 min in PBS-T. Finally, samples were mounted in Vectashield, and NMJs were imaged using a Zeiss LSM800 confocal microscope. The number of postsynaptic boutons and branches was manually quantified from five different larvae per genotype analyzing three NMJs corresponding to muscle 12 in segments A2-A4, with at least one NMJ from both the right and left hemisegments.

### Fertility Assay

The fertility assay was performed by quantif ying the laid eggs from fertilized females of different genotypes. For that, two crosses were set up with ten 1- to 3-day-old control males and ten 1- to 3-day-old control (*w^1118^;;*)or *dwdr45* KO (*w^1118^;;dwdr45KO/TM6B,Tb*) virgin females for three days at 25°C. Flies were then transferred to a fertility assay chamber with laying medium (300 mL H2O, 100 mL grape juice, 10 g sucrose [Euromedex, 200-301-B], 9 g agar [VWR International, 20768.235], 0.6 g nipagine [Sigma-Aldrich, H3647]). Twice per day and for five days, laid eggs were counted, and a new medium was provided. After 48 h, the number of unhatched eggs was counted: yellow eggs were considered unfertilized, dark eggs were considered as eggs with embryonic development disturbing. The number of hatched eggs corresponds to the difference between laid eggs and unhatched eggs.

### Lifespan Analysis

For lifespan analysis, 6 vials of 20 adult females per condition were harvested within one day of adult hatching and maintained at 25°C. Every two or three days, flies were transferred to a new vial and longevity was scored by counting the number of dead flies. Flies that escaped from the vial were recorded as censored. The Kaplan-Meier curve represents the percentage of surviving flies depending on time. Statistical analyses were performed on R Software using ‘survival’ package. Cox test was performed to allow for comparison of multiple conditions based on hazard ratio that corresponds to the probability of fly death event apparition.

### Western Blot Analysis

Fly heads (n=10) were mixed in 50 µL of Laemmli buffer 1X (Sigma-Aldrich, S3401) and boiled for 5 min before freezing. Proteins were separated by sodium dodecyl sulfate-polyacrylamide gel electrophoresis (BIO-RAD, 4568093) at 100 V during 1 h before transferring on polyvinylidene difluoride membrane (Thermo Fisher Scientific, 88518) at 25 V overnight at 4°C. Membrane was washed three times for 5 min each in Tris-Buffered Saline (TBS) + 0.1% Tween-20 (Sigma-Aldrich, P1379) before 10 min saturation in blocking solution (TBS1X + 0.1% Tween-20 + 5% BSA). The primary antibodies incubation was performed overnight at 4°C with mouse anti-α-Tubulin (1:2000 [Sigma-Aldrich, T6199]), rabbit anti-Gabarap (1:2000 [abcam, ab109364]), rabbit anti-Ref(2)P (1:1000 [abcam, ab178440]), rabbit anti-Fer1HCH (1:500) or rabbit anti-Fer2LCH (1:500) (Gift from Fanis Missirlis, (60)) diluted in blocking solution. The membrane was washed three times for 5 min before secondary antibodies incubation 1 h at room temperature with anti-Mouse-A488 (1:10 000 [Thermo Fisher Scientific, A-11001]) and anti-Rabbit-A546 (1:10 000 [Thermo Fisher Scientific, A-11035]) diluted in blocking solution. The membrane was finally washed three times for 5 min each before fluorescence monitoring using ChemiDoc MP Imaging System (Bio-RAD). Analysis was realized by quantification of band fluorescence intensity using ImageLab software (BIO-RAD).

### Ferrozine Assay

Ferrozine assays were adapted from Riemer et al., (61). Briefly, the whole bodies of 3 flies were ground in 125 µL of 50mM NaOH and kept for 2 h at 4°C on a rotor. 100 µL of lysate and 100 µL of 10 mM HCl were mixed before adding 100 µL of the iron-releasing reagent (50 µL of 1.4 M HCl + 50 µL of 4.5% KMnO4 [Sigma-Aldrich, 223468] diluted in H2O). Samples were heated 2 h at 60°C before cooling at room temperature. Finally, 20 µL of iron-detection solution (6.5 mM ferrozine [Sigma-Aldrich, 82940], 6.5 mM neocuproine [Sigma-Aldrich, 72090], 2.5 mM ammonium acetate [Sigma-Aldrich, A1542], 1 M ascorbic acid [Sigma-Aldrich, A5960] diluted in H2O) were added. Absorbance was read in triplicate on a 96-well plate at 560 nm. The iron concentration was determined by using a standard curve and normalized against the protein concentration measured using the Bradford assay on 5 µL of lysate. For each condition, quadruplicates were performed.

### Whole mount adult retina immunostaining

For each condition, 8 retinae of 8-day-old adult female flies were dissected in Schneider medium and stored in Schneider + 1% PFA on ice until end of dissections. Brains were fixed in Schneider + 4% PFA for 20 min at room temperature followed by three-times 15 min washes in washing solution (PBS1X + 0.3% Triton-X-100 + 5 mg/mL BSA). Saturation was realized over 30 min in blocking solution (washing solution + 4% NGS). Retinae were incubated in primary antibodies overnight at 4°C with rabbit anti-GFP (1:200 [Thermo Fisher Scientific, A6455]) or rabbit anti-Ref(2)P (1:400 [provided by S. Gaumer]) diluted in blocking solution. Retinae were washed three times for 15 min each before secondary antibodies incubation for 2 h at room temperature with anti-Rabbit-A488 and Phalloidin-Rhodamine (1:400 [Thermo Fisher Scientific, R415]) diluted in blocking solution. Finally, retinae were washed three times for 15 min each mounted in Vectashield on bridge slides. Confocal microscopy was done using a Zeiss LSM800 microscope.

Quantification of fluorescent percentage area was performed in ImageJ by segmenting puncta using uniform threshold parameters across all samples. The total puncta area was then computed for each image and normalized to the corresponding total retinal area to obtain the percentage of fluorescent coverage.

### Adult retina immunostaining after horizontal cryosectioning

For each condition, 6 fly heads were cut in PBS1X and placed in OCT (Electron Microscopy Sciences, 62550) before 1h freezing in dry ice. 10 µm slices were cut using HM525 NX Cryostat and collected on superfrost slides (VWR International, 48404-476). Slides were dried 15 min before fixation in PBS1X + 4% PFA for 1 h at room temperature. Slides were briefly washed in three successive PBS1X baths before saturating in blocking solution (PBS1X + 0.3% Triton-X-100 + 1% BSA) over 30 min. Primary antibody incubation was realized for 1h at room temperature with rabbit anti-GFP (1:400) or rabbit anti-Ref(2)P (1:400 [provided from S. Gaumer]) diluted in blocking solution. Slides were briefly washed in three successive PBS1X baths before secondary antibody incubation for 1 h with anti-rabbit-A647 (1:400 [Thermo Fisher Scientific, A-31573]) diluted in blocking solution. Finally, slides were briefly washed in three successive PBS1X baths before mounting in Vectashield with DAPI (Eurobio Scientific, H-1200). Confocal microscopy was performed using a Zeiss LSM800 microscope.

Quantification of Ref(2)P dots was performed in ImageJ by segmenting puncta using uniform threshold parameters across all samples. Puncta number was subsequently quantified using the *Analyze Particles* function. In conditions of Ref(2)P::GFP expression, threshold-based segmentation did not allow clear puncta isolation. Therefore, mean fluorescence intensity within the retina was measured instead.

### Statistical Analyses

Statistical analyses were performed in JAMOVI, using R software. Samples were compared through one-way or two-way ANOVA depending on the number of parameters. In addition, when ANOVA indicates a significant difference, post-hoc tests were performed using different corrections depending on variance’s homogeneity: Tukey’s correction for homoscedasticity or Games-Howell’s correction for heteroscedasticity. On graphs, data are reported as the means ± standard error of mean (SEM) of three experiments unless otherwise noted.

#### Ethic statement

All experiments were realized using *Drosophila melanogaster* with the authorization for the contained use of class 2 genetically modified organisms from the French Ministry of Higher Education and Research (Duo No. 11960).

## Acknowledgments

We are very grateful to the patient associations and funders who supported this work. We thank our colleagues from the fly community and fly stock centers for providing fly stocks and antibodies used in this study. We thank B. Horard, LBMC, for her help with fertility experiments. We thank the imaging LYMIC-PLATIM and the fly ARTHRO-TOOLS facilities (SFR Biosciences, Lyon). Chat GPT 3.5. and Perplexity AI were used for English language editing in some sections of this manuscript. Biorender was used to generate the graphical abstract. This article is dedicated to Dr Francine Côté, who always supported the fly as a great model to advance BPAN research.

## Abbreviations

ATG: autophagy-related
BPAN: beta-propeller protein-associated neurodegeneration
KD: knock-down
KO: knock-out
NBIA: neurodegeneration with brain iron accumulation
NMJ: neuromuscular junction
WDR45: WD repeat domain 45
WIPI: WD40 repeat protein interacting with phosphoinositides.

