## Supplemental Figures for "A *dwdr45* knock-out *Drosophila* model to decipher the role of autophagy in BPAN"

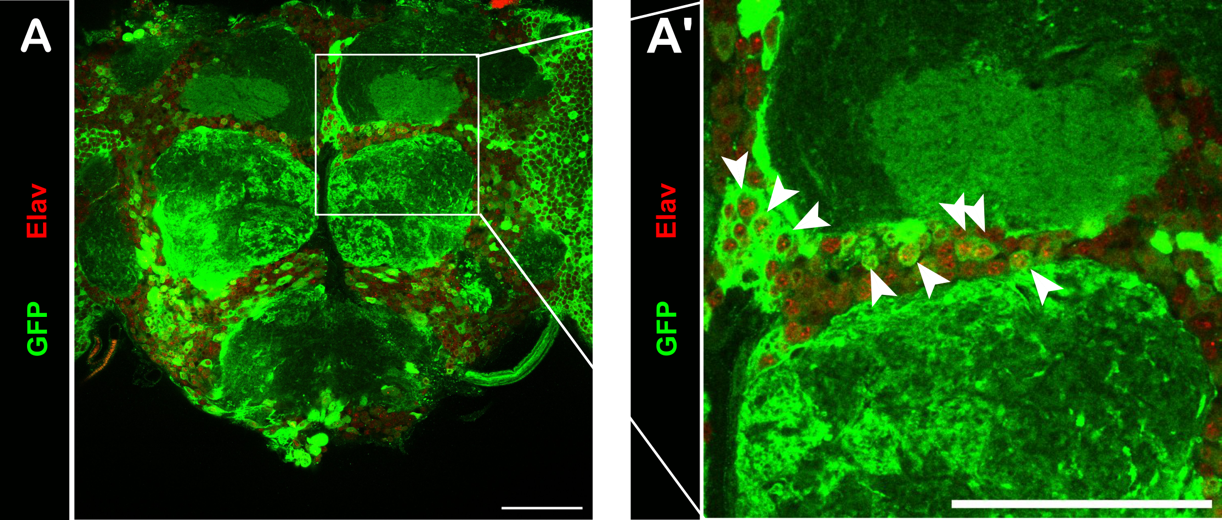

**Figure S1. Expression of *dWdr45* in *Drosophila* adult brain.**

*dWdr45* expression in brain is shown using the GAL4 *CG11975^KO-kG4^* promoter trap line combined with UAS-GFP *(UAS-GFP/+ ; CG11975^KO-kG4^* */+*). GFP Immunostaining and neuronal colabelling were realized on brain dissection of 8-days-old adult flies using an anti-GFP and anti-elav antibodies. Arrow heads show intense neuronal expression of GFP in the anterior brain. Scale bars represent 50 μm.

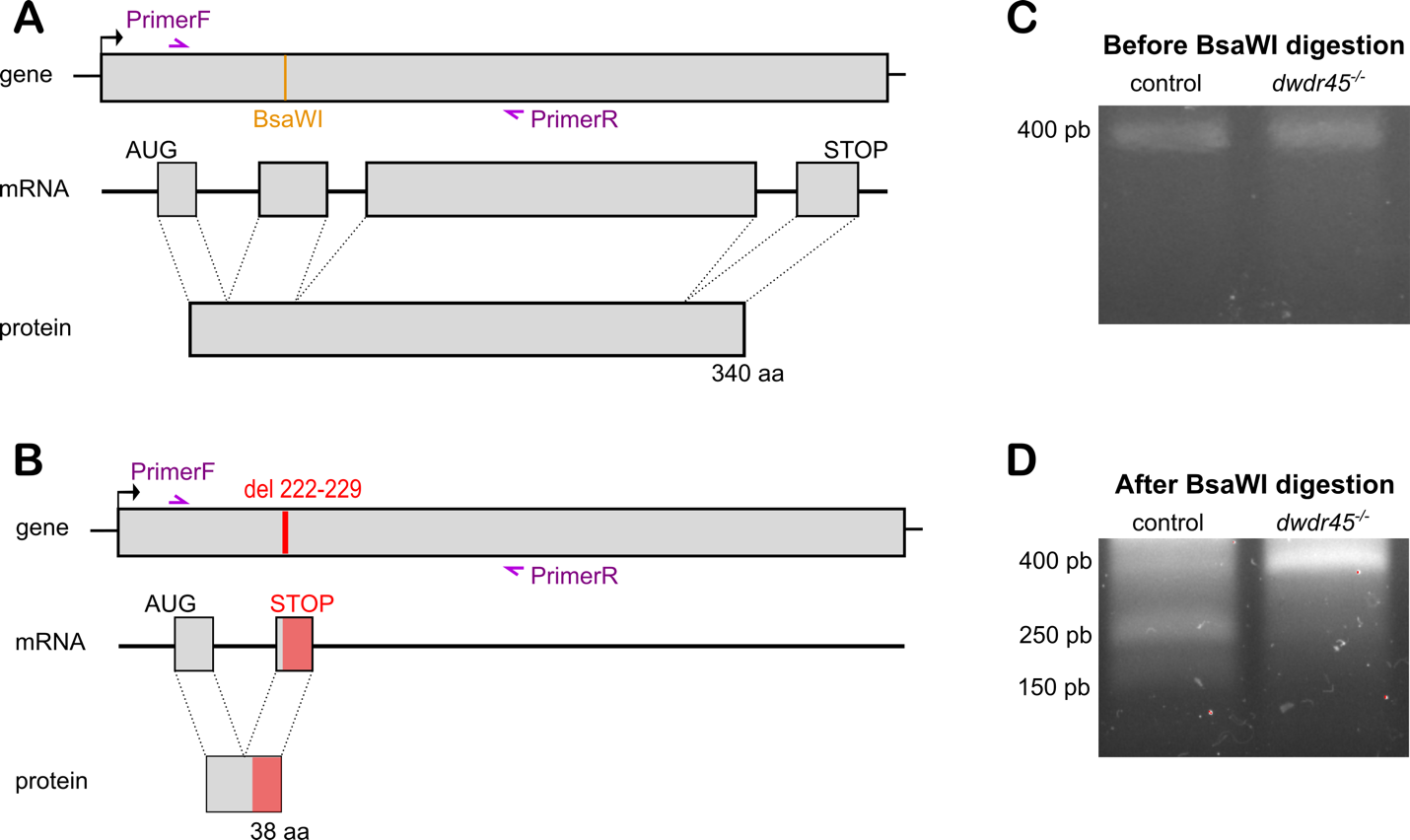

**Figure S2: Crispr/CAS9-mediated KO of dWdr45 gene.**

(**A**) Schematic representation of wild type *dWdr45* gene, mRNA and protein. Primers used for genotyping are show in purple and BsaWI digestion site in yellow. (**B**) Schematic representation of *dwdr45* KO gene after Crispr/CAS9-mediated deletion of seven nucleotides including BsaWI digestion site. This deletion led to a frameshift and a premature stop codon during the translation. (**C**) Electrophoresis gel after DNA amplification using described primers. Both control and homozygous *dwdr45* KO flies show a 397pb fragment. (**D**) Electrophoresis gel of DNA amplification digest by BsaWI. In control flies, BsaWI recognized digestion site and cut DNA leading to a 153pb and a 254pb fragment. In homozygous *dwdr45* KO flies, DNA is not cut due to the absence of BsaWI digestion site.

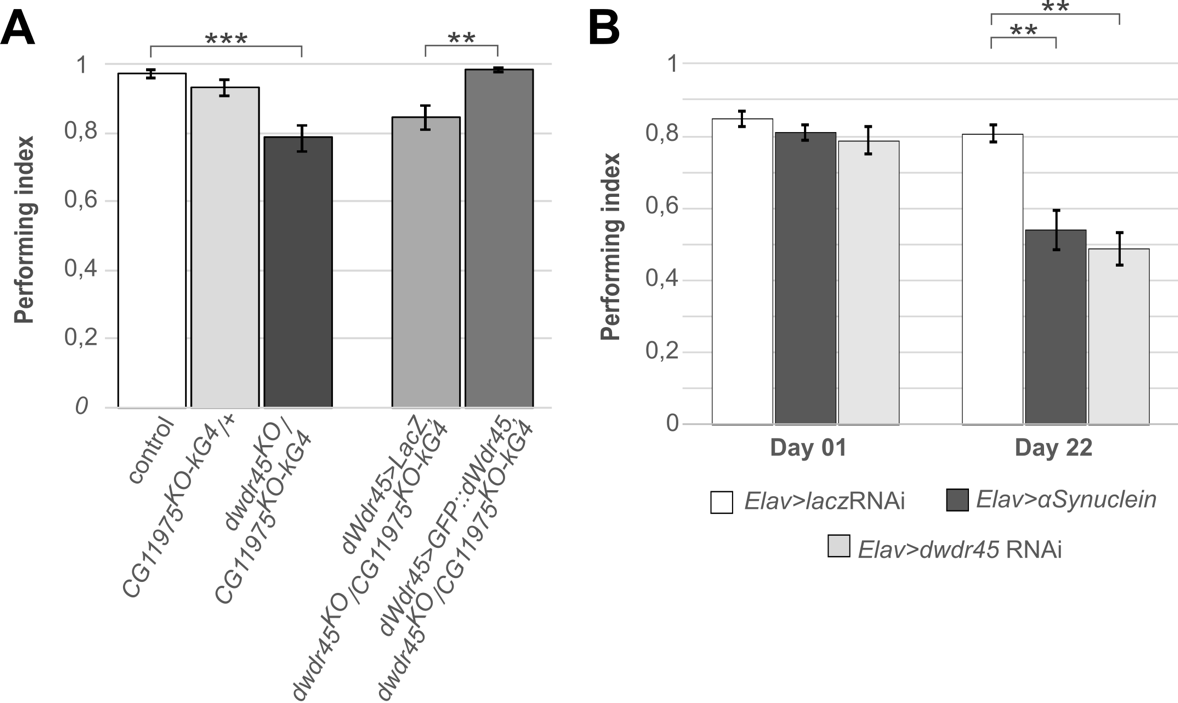

**Figure S3: Locomotor disorder in BPAN *Drosophila* model**

(**A**) SING assays were realized in 8-day-old adult control, heterozygous *CG11975^KO-kG4^* or transheterozygous *CG11975^KO-kG4^ /dwdr45*^KO^ flies. Overexpression of LacZ or GFP::dWdr45 is also realized in transheterozygous flies under dWdr45 driver. Statistical analyses were carried out in three independent experiments (n=6) with ANOVA2/post-hoc test: **p-value<0.01 and ***p-value<0,001. (**B**) SING assays were realized in 8-day-old adult flies expressing *lacz* RNAi, α-synuclein or *dwdr45* RNAi under the pan-neuronal Elav driver. Statistical analyses were carried out in three independent experiments (n=6) with ANOVA2/post-hoc test: **p-value<0.01.

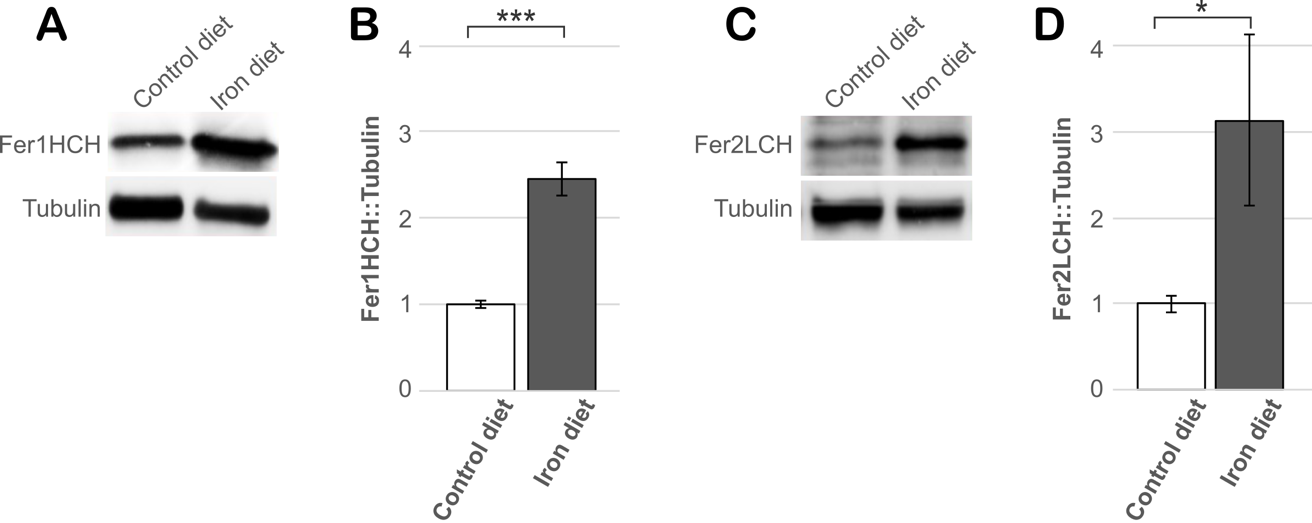

**Figure S4: Ferritins levels are sensitive to iron overload.**

(A) Fer1HCH proteins are detected by western blotting in 4-day-old adult wild-type flies fed with a control diet or an iron-supplemented diet (30mM FAC). (B) The ratio of Fer1HCH relative to tubulin protein levels is normalized to control diet. Statistical analyses were carried out in four independent experiments with ANOVA2/post-hoc test: ns: no-significant, ***p-value<0.001. (C) Fer2LCH proteins are detected by western blotting in 4-day-old adult wild-type flies fed with a control diet or an iron-supplemented diet (30mM FAC). (D) The ratio of Fer2LCH relative to tubulin protein levels is normalized to control diet. Statistical analyses were carried out in four independent experiments with ANOVA2/post-hoc test: ns: no-significant, *p-value<0.05.

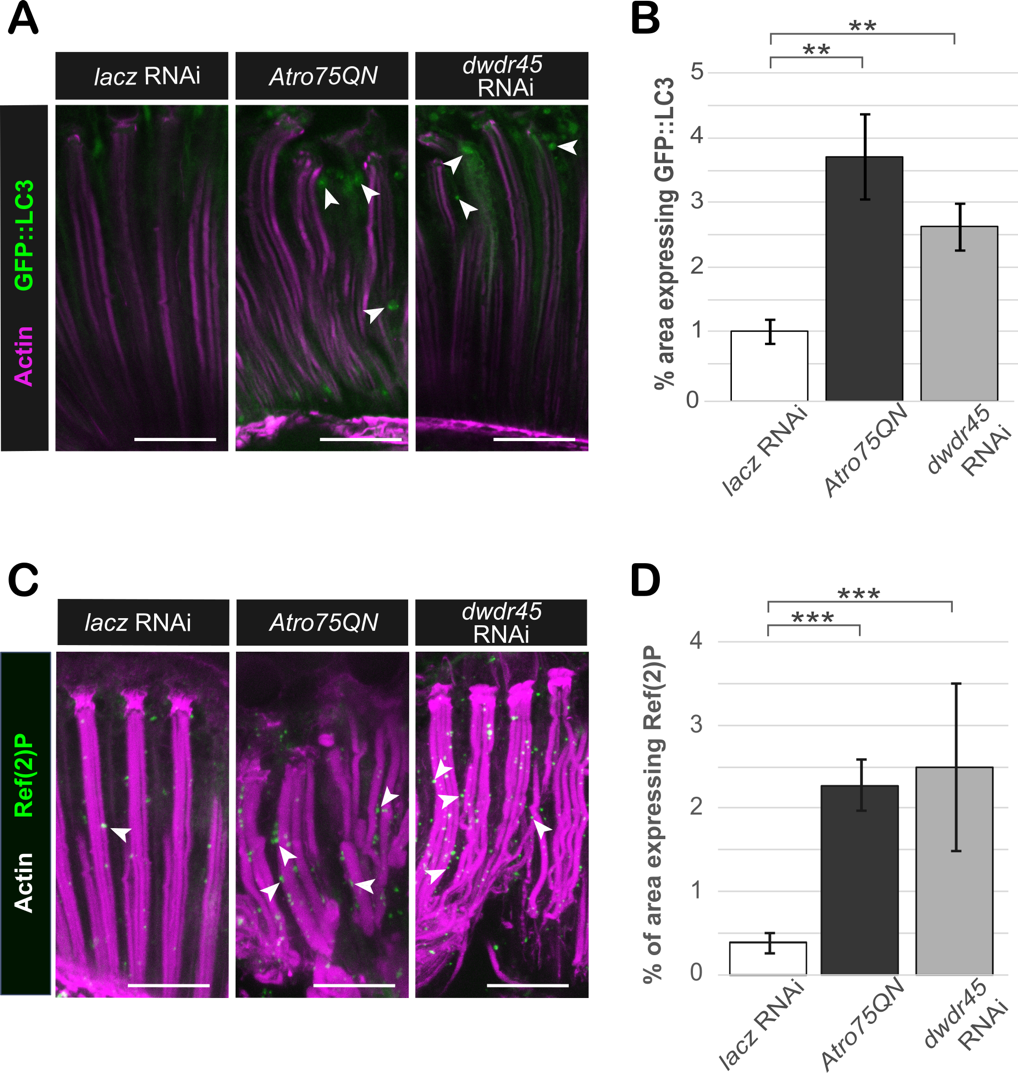

**Figure S5: Retinal accumulation of GFP-LC3 and Ref(2)P in pan-retinal KD of *dwdr45*.**

(**A**) Immunolabelling of GFP in dissected adult retina that overexpress *GFP::LC3* under the pan-retinal GMR driver in 22-day-old adult flies also expressing *lacz* RNA, *Atro75QN* or *dwdr45* RNAi. Phalloïdin was used to label actin. Scale bars represent 20µm. (**B**) Quantification of the percentage of retina area expressing GFP. Statistical analyses were carried out in three independent experiments (n=6) with ANOVA1/post-hoc test: **p-value<0.01. (**C**) Immunolabelling of Ref(2)P protein in dissected retina of 22-day-old adult flies expressing *lacz* RNAi, *Atro75QN* or *dwdr45* RNAi under the pan-retinal GMR driver. Phalloïdin was used to label actin. Scale bars represent 20 µm. (**D**) Quantification of the percentage of retina area expressing *Ref(2)P*. Statistical analyses were carried out in three independent experiments (n=6) with ANOVA1/post-hoc test: **p-value<0.01.
